# A refined InDel taxonomy broadens the repertoire of mutational processes shaping multiple myeloma

**DOI:** 10.64898/2026.11.23.768572

**Authors:** Marios Papadimitriou, Abhishek Pandey, Alexandra Poos, Nathan Becker, Abdessamad Youssfi Alaoui, Goren Arik, Nicholas Sangvai, Michael Durante, Chrissy Baker, Pedro Manuel Davila, Enze Liu, Sreejeta Mondal, Tomas Jelinek, Bachisio Ziccheddu, Francesco Maura, Florence Magrangeas, Stéphane Minvielle, Elias Karl Mai, Katja Weisel, Roland Fenk, Marc-Steffen Raab, Hartmut Goldschmidt, Ola Landgren, Eric Letouze, Brian Walker, Niels Weinhold, Benjamin Diamond

## Abstract

Somatic aberrations have been extensively characterized in multiple myeloma (MM). However, the mutational processes underlying small insertions and deletions (InDels) remain poorly understood. Here, we used a recently developed 89-channel InDel classification schema which allows differentiating mutational processes. We analyzed whole-genome sequencing data from 887 MM samples, spanning both newly diagnosed (NDMM) and relapsed/refractory disease (RRMM). We identified InDel signatures associated with replication slippage, APOBEC mutagenesis, topoisomerase 1 (TOP1)-mediated mutagenesis, non-homologous end joining (NHEJ), and two rare DNA damage repair-deficiency signatures. In two randomized clinical trials, incorporation of the APOBEC InDel signature InD9a improved treatment outcome prediction as pertaining to progression-free survival (PFS) and achievement of minimal residual disease (MRD) negativity. InD29, a signature linked to NHEJ-mediated errors, was detected in both NDMM and RRMM, and was enriched in samples exhibiting melphalan-associated mutagenesis and high SV burden. Genome-wide analysis identified a recurrent InDel hotspot at *TXNDC5*, a highly expressed gene in MM located within a region implicated in complex genomic rearrangements, with predominantly noncoding alterations suggesting potential regulatory consequences. Finally, InDel signatures revealed the presence of mismatch-repair deficiency (MMRd) and homologous recombination deficiency (HRD) phenotypes, even in cases without germline predisposition. Together, these findings provide a comprehensive characterization of InDel mutagenesis in MM, revealing previously underexplored relationships and demonstrating potential to refine comprehensive genomic classifications of MM.

**KEY POINTS:**

- InDel mutational signatures reveal distinct mutational processes in multiple myeloma and improve prognostic stratification beyond SBS signatures.
- InDel mutational signatures identify rare DNA repair-deficient multiple myeloma subgroups characterized by hyperhaploidy.

## INTRODUCTION

Multiple myeloma (MM) is a plasma cell malignancy characterized by extensive genomic heterogeneity, arising through the acquisition of recurrent copy number alterations (CNVs), somatic single nucleotide variants (SNVs), and complex structural variants (SVs)^1–8^. Whole-genome sequencing (WGS) studies have demonstrated that MM evolution begins decades before clinical diagnosis, with canonical immunoglobulin heavy chain (*IGH*) translocations and hyperdiploidy representing early genomic events, followed by the acquisition of subsequent driver alterations that contribute to disease progression^4,9–13^. The characterization of mutational signatures has further enabled the identification of the biological processes responsible for generating these alterations throughout tumor evolution^3,4,14–17^.

Single-base substitution (SBS) signatures have enabled the characterization of distinct mutational processes that contribute to punctuated and ongoing mutagenesis and clonal diversity^9,11,15,18–23^. Copy number signatures have been associated with mechanisms of chromosomal instability (ie chromothripsis)^24–26^. Similarly, SV signatures have revealed distinct patterns of rearrangements and recurrent SV hotspots implicated in MM pathogenesis^27–29^. In contrast, small insertions and deletions (InDels) have received comparatively less attention, partly owing to the analytical complexity associated with their detection and classification^1,30,31^. Large-scale pan-cancer analyses have established the Catalogue of Somatic Mutations in Cancer (COSMIC) InDel signature framework^15^. This classification schema has been used to investigate therapy-related InDel mutational signatures in hematologic malignancies, including MM, identifying the non-homologous end joining (NHEJ) signature ID8 in relapsed hematologic malignancies following exposure to radiotherapy or mutagenic chemotherapy^32^. However, low resolution in key channels, such as homopolymer or small (1 base pair; bp) InDel channels have limited further discovery.

Recently, Koh et al. (2025) introduced a redefined 89-channel InDel classification incorporating flanking sequence context, homopolymer tract lengths, and repetitive sequence motifs, substantially improving the discrimination of biologically distinct mutational processes^33^. Building upon this framework, we comprehensively characterized the landscape of somatic InDels in MM using WGS data from 887 samples. Through *de novo* mutational signature extraction and fitting, we systematically investigated the biological processes underlying InDel mutagenesis and their contributions to genomic and clinical heterogeneity. Overall, we identify an APOBEC InDel signature present in MM that adds prognostic resolution to the existing SBS signatures, identify NHEJ as a prominent process associated with SV accumulation and melphalan mutagenesis, and describe two novel DNA damage repair-deficient MM subgroups, each with therapeutic implications. Our study provides a comprehensive characterization of the InDel mutational signature landscape in MM, revealing previously underappreciated associations and MM subtypes with clinical relevance.

## METHODS

### Study cohorts

We analyzed WGS data from 887 samples; this included a discovery cohort comprised of 466 samples and an independent validation cohort with 421 samples. The discovery cohort comprised 466 samples from 421 patients, including 319 NDMM and 147 RRMM. Of these, 378 patients contributed a single sample, 41 patients contributed 2 samples, and 2 patients contributed 3 samples. These samples originated from three institutions: Heidelberg University (n = 360 samples; 322 patients)^34,35^, Indiana University [Indiana Myeloma Registry (IMR); n = 52 samples; 52 patients], and Sylvester Comprehensive Cancer Center (SCCC; n = 54 samples; 47 patients). For the Heidelberg University cohort, 264 NDMM samples were collected as part of the GMMG-HD6 trial and 96 RRMM samples were collected independently of the trial. The validation cohort consisted of 421 NDMM patients (n = 421 samples) from the MIDAS (IFM) trial^36,37^. This study design enabled the characterization of somatic InDel patterns across disease settings and provided an independent cohort to assess the reproducible extraction of the identified signatures. Sample collection and processing procedures differed across the three institutions, as detailed in the **Supplemental Methods**. Tumor and germline control samples were collected under the respective Institutional Review Boards’ protocols for all four included cohorts, with written informed consent for all participants, and in accordance with the Declaration of Helsinki.

### Mutational signature analysis

Mutational signatures were analyzed across all MM datasets. For SBS signatures, we used the R package *mmsig* (https://github.com/UM-Myeloma-Genomics/mmsig) and fitted the established group of SBS signatures known to show activity in MM^9,38^. To assign the *de novo* extracted InDel signatures to established mutational processes, we performed pairwise fitting against the reference InDel signatures described by Koh et al., 2025^33^. For each *de novo* signature, all possible two-signature combinations were evaluated, and their relative contributions and cosine similarities (cossim) were calculated to identify the reference signatures providing the best reconstruction (https://github.com/UM-Myeloma-Genomics/Signature-Assignment). Subsequently, reference InDel signatures were fitted to individual samples using a modified version of *mmsig*. Details of the sequencing pipeline as well as further computational analyses are provided in the **Supplementary Methods**.

## RESULTS

### The InDel landscape of multiple myeloma

To characterize the landscape of somatic InDels in MM, we analyzed WGS data from 887 samples; this included a discovery cohort comprised of 466 samples and an independent validation cohort with 421 samples (**Figure 1A**; n = 887). Across the discovery set, we identified a total of 239,256 somatic InDels, including 142,448 in NDMM and 96,808 in RRMM (**Figure 1B**). Examination of the InDel profiles across the cohorts demonstrated largely comparable distributions of mutational classes despite differences in the InDel calling pipelines employed (**Supplementary Figure 1**). Comparison of NDMM and RRMM revealed a significantly higher per-sample InDel count in RRMM (mean = 659; range, 158–1,853) compared to NDMM (mean = 447; range, 55–8,257; Wilcoxon p = 5.05 × 10^-19^), consistent with previous reports of increased mutational burden in advanced MM (**Figure 1C**)^3,11,39^. We next investigated whether specific InDel classes contributed disproportionately to the increased mutational burden observed in RRMM. Single-base cytosine (1-bp C) deletions (Wilcoxon p < 0.001; **Figure 1D**) and microhomology (Mh)-associated deletions (Wilcoxon p < 0.001; **Figure 1E**) were significantly enriched in RRMM compared to NDMM. These differences were also reflected in their relative contributions to the overall InDel repertoire, with 1-bp C deletions increasing from 9.6% in NDMM to 11.2% in RRMM (Wilcoxon p < 0.001), and Mh deletions slightly increasing from 8.9% to 9.2% (Wilcoxon p = 0.023). Conversely, the relative contribution of single-base thymine (1-bp T) deletions decreased from 29.4% in NDMM to 26.2% in RRMM (Wilcoxon p < 0.001; **Supplementary Figure 2; Supplementary Tables 1-2**). Together, these findings indicate non-random enrichment of specific deletion classes across the disease continuum, suggesting an increase in mutational activity of discrete processes^40–42^.

**Figure 1.**
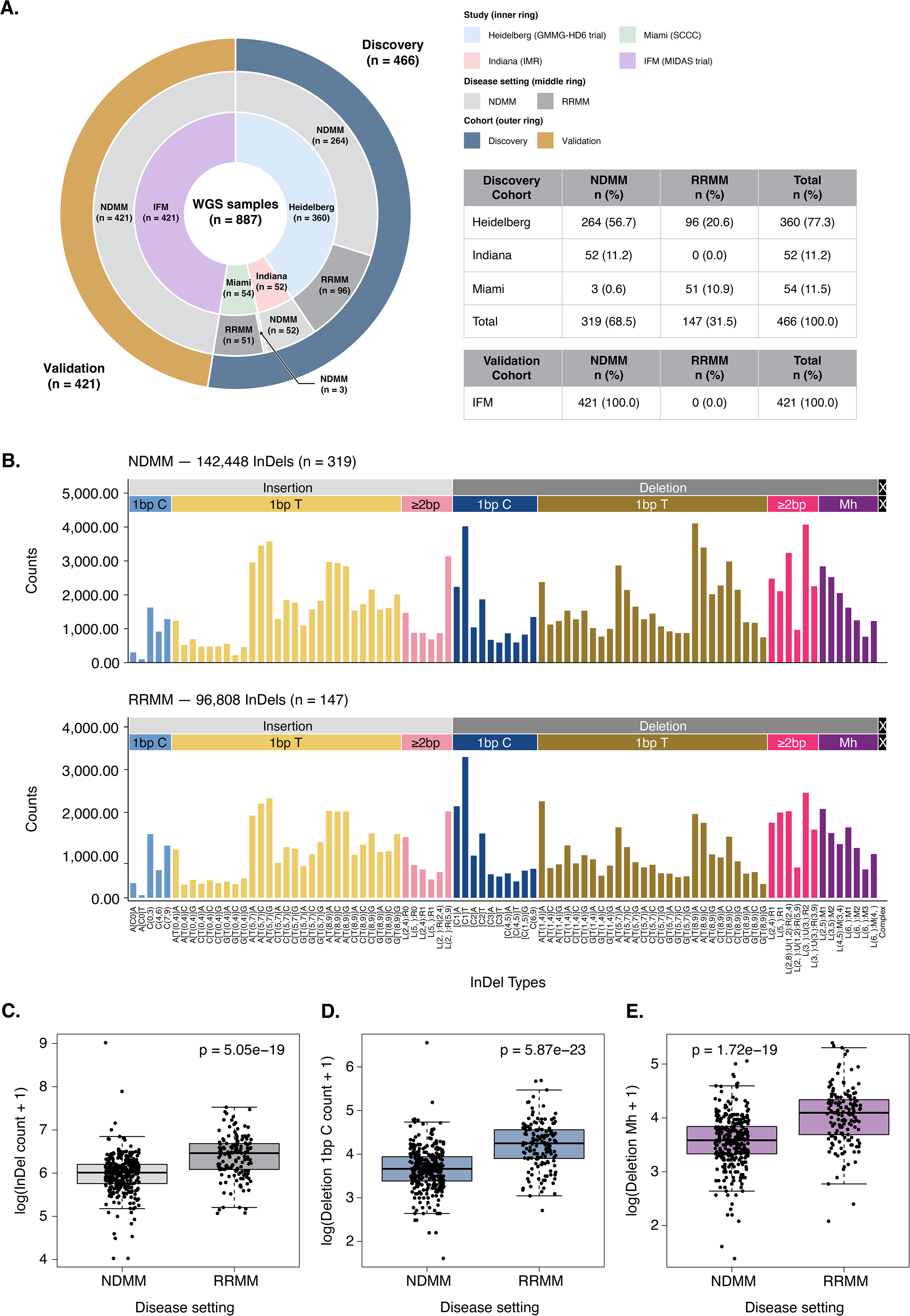
The InDel landscape of MM. **(A)** Overview of the study cohorts comprising 887 WGS samples, including 466 samples in the discovery cohort from Heidelberg (GMMG-HD6; n = 360), Indiana (IMR; n = 52), and Miami (SCCC; n = 54), and 421 samples in the independent validation cohort from the IFM (MIDAS trial). The concentric rings indicate study of origin (inner), disease setting (middle), and cohort assignment (outer). The accompanying tables summarize sample distributions across NDMM and RRMM within the discovery and validation cohorts, respectively. **(B)** Distribution of 239,256 somatic InDels across the 89-channel InDel classification framework, stratified by disease setting (NDMM, n = 319 samples, 142,448 InDels; RRMM, n = 147 samples, 96,808 InDels). InDels are grouped according to type, nucleotide context, repeat length, and microhomology characteristics. Bar heights represent the aggregate number of InDels assigned to each mutational channel. **(C)** Comparison of total InDel burden, **(D)** 1-bp cytosine deletion burden, and **(E)** Mh-associated deletion burden between NDMM and RRMM samples in the discovery cohort. Values are log-transformed as log10(count + 1). Each point represents an individual sample, and boxes indicate the interquartile range with the central line representing the median. P values are displayed in each panel. MM: multiple myeloma; WGS: whole-genome sequencing; NDMM: newly-diagnosed multiple myeloma; RRMM: relapsed/refractory multiple myeloma; InDel: insertion and deletion; bp: base pair; Mh: microhomology.

### De novo extraction and deconvolution of InDel mutational signatures

To identify the underlying InDel mutational signatures responsible for InDel burden, we applied the *de novo* signature extraction framework as described by Koh et al.^33^ to the discovery cohort (n = 466). Briefly, the algorithm employs non-negative matrix factorization (NMF) to decompose the observed mutational catalogue into distinct signatures and quantifies their corresponding contributions across individual samples. A four-signature solution provided the most favorable balance between reconstruction accuracy and stability and was therefore selected for subsequent analyses (**Supplementary Figure 3**). The four extracted signatures exhibited distinct non-overlapping mutational profiles (**Figure 2A**, **Supplementary Table 3**). SIG_1 was predominantly characterized by 1-bp T insertions at homopolymer tracts of varying lengths, alongside a prominent contribution from ≥2-bp insertions. SIG_2 was enriched in ≥2-bp deletions, particularly those involving microhomology (Mh) at deletion junctions. SIG_3 exhibited prominent peaks of 1-bp C deletions, particularly at short homopolymer tracts, accompanied by ≥2-bp deletions with limited Mh deletions. Finally, SIG_4 was dominated by 1-bp T deletions at extended homopolymer tracts, particularly those comprising 8–9 nucleotides, with additional contributions from deletions at shorter repeats (5–7 nucleotides).

**Figure 2.**
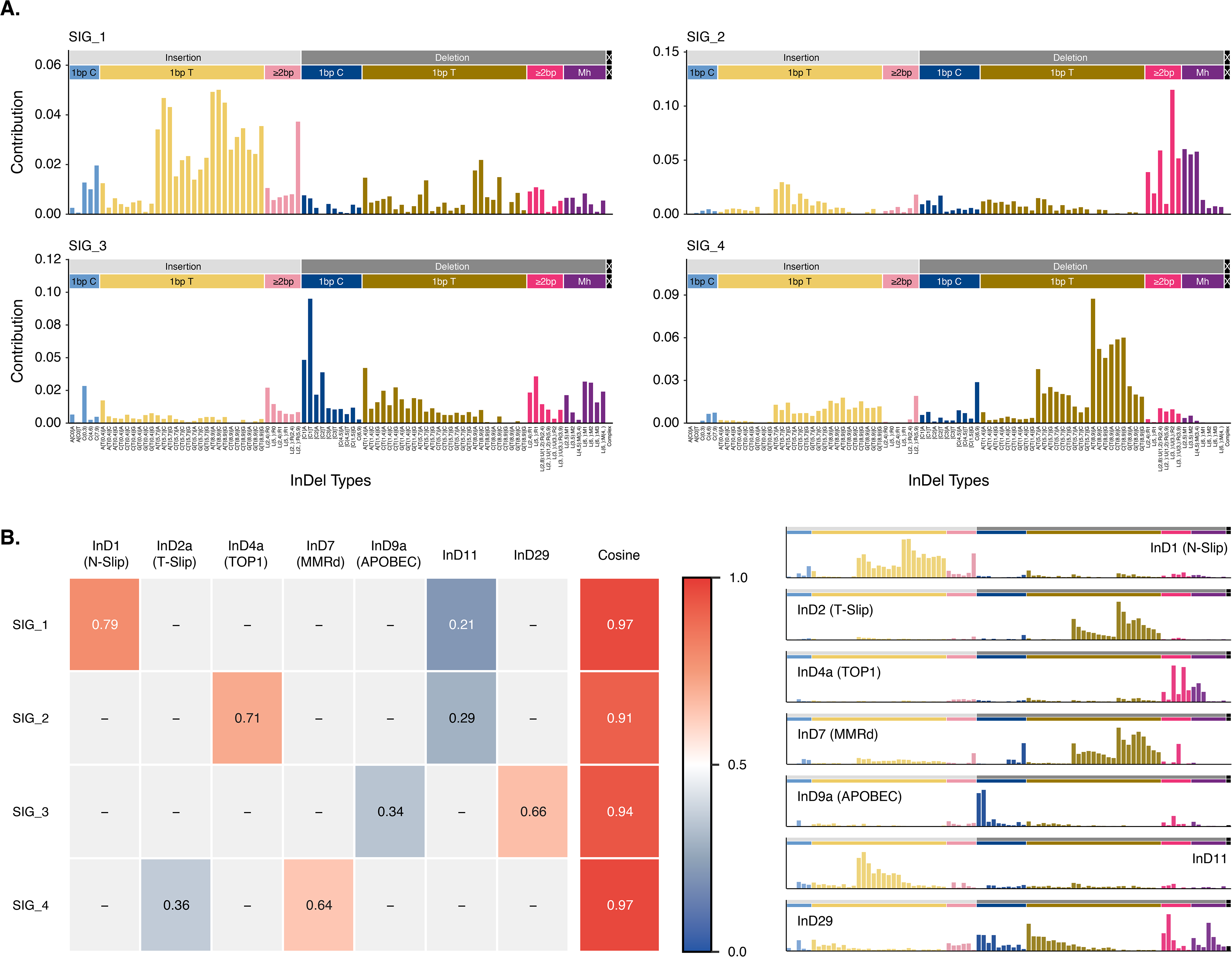
*De novo* extraction and deconvolution of InDel mutational signatures in MM. **(A)** Mutational profiles of the four *de novo* InDel signatures (SIG_1–SIG_4) extracted from the discovery cohort (n = 466 samples) using NMF and the 89-channel InDel classification framework. Bar heights represent the relative contribution of each mutational channel to the corresponding signature. **(B)** Decomposition of the four *de novo* signatures into seven previously described reference InDel signatures (reference). The heatmap displays the relative contribution of each reference signature to the reconstructed *de novo* signatures, with cossim values indicating the agreement between the original and reconstructed profiles. The mutational profiles of the seven reference signatures are shown on the right. Reference signatures include InD1 (N-Slip), InD2a (T-Slip), InD4a (TOP1), InD7 (MMRd), InD9a (APOBEC), InD11, and InD29. NMF: non-negative matrix factorization; cossim: cosine similarity; N-Slip: Nascent-strand slippage; T-Slip: Template-strand slippage; TOP1: DNA topoisomerase 1; MMRd: mismatch repair deficiency.

To relate the extracted signatures to known mutational processes, we performed pairwise deconvolution of each *de novo* signature into a combination of two reference signatures as previously described (**Figure 2B**; **Supplementary Tables 4-7, Methods**)^23,32,33^. The resulting decompositions were able to reconstruct the *de novo* extracted signature achieving high cosine similarity (cossim > 0.90; **Figure 2B**)^33^. The identified reference signatures reflected several distinct endogenous mutational processes, some of which parallel previously characterized SBS signatures in MM. InD1 and InD2a have been attributed to slippage during DNA replication, reflecting replication-associated mutagenesis common to many malignancies. InD4a has been linked to TOP1-associated mutagenesis, whereas InD9a has been associated with APOBEC activity and correlated with SBS2/SBS13, two well-characterized mutational signatures in MM^40,42^. Notably, InD29 shares features with InD8, a signature reflecting NHEJ activity, although whether InD29 represents a related or distinct mutational process remains unclear^33^. InD11 shares features with InD1 and may represent a similar process. Finally, we identified InD7, a signature associated with MMRd, only recently described in MM^37^.

### InD9a is linked to APOBEC mutagenesis and adds prognostic resolution

To assess the sample-level contribution of the deconvoluted mutational signatures, we developed a modified version of *mmsig*, adapted to incorporate the 89-channel InDel mutational classification, and applied it to the discovery cohort for signature fitting (**Figure 3A**, **Methods, Supplementary Table 8**). Given the extensive characterization of APOBEC (SBS2/SBS13) in NDMM and its established clinical relevance^6,9,20,21,43^, we next focused on the InDel signature InD9a, proposed as related to APOBEC mutagenesis^33^. Although their relative contributions were positively correlated (Spearman’s ρ = 0.4, p < 0.001), we observed an interesting finding; notably, there was a group of samples with low SBS2/SBS13 contribution but substantial InD9a contribution (**Supplementary Figure 4A**)^6,43^. To ascertain whether this phenomenon was specific to MM or our cohort, we assessed SBS2/SBS13 and InD9a in breast tumors from the BRCA-EU cancer cohort, and observed a similar relationship (**Supplementary Figure 4B**)^44,45^.

**Figure 3.**
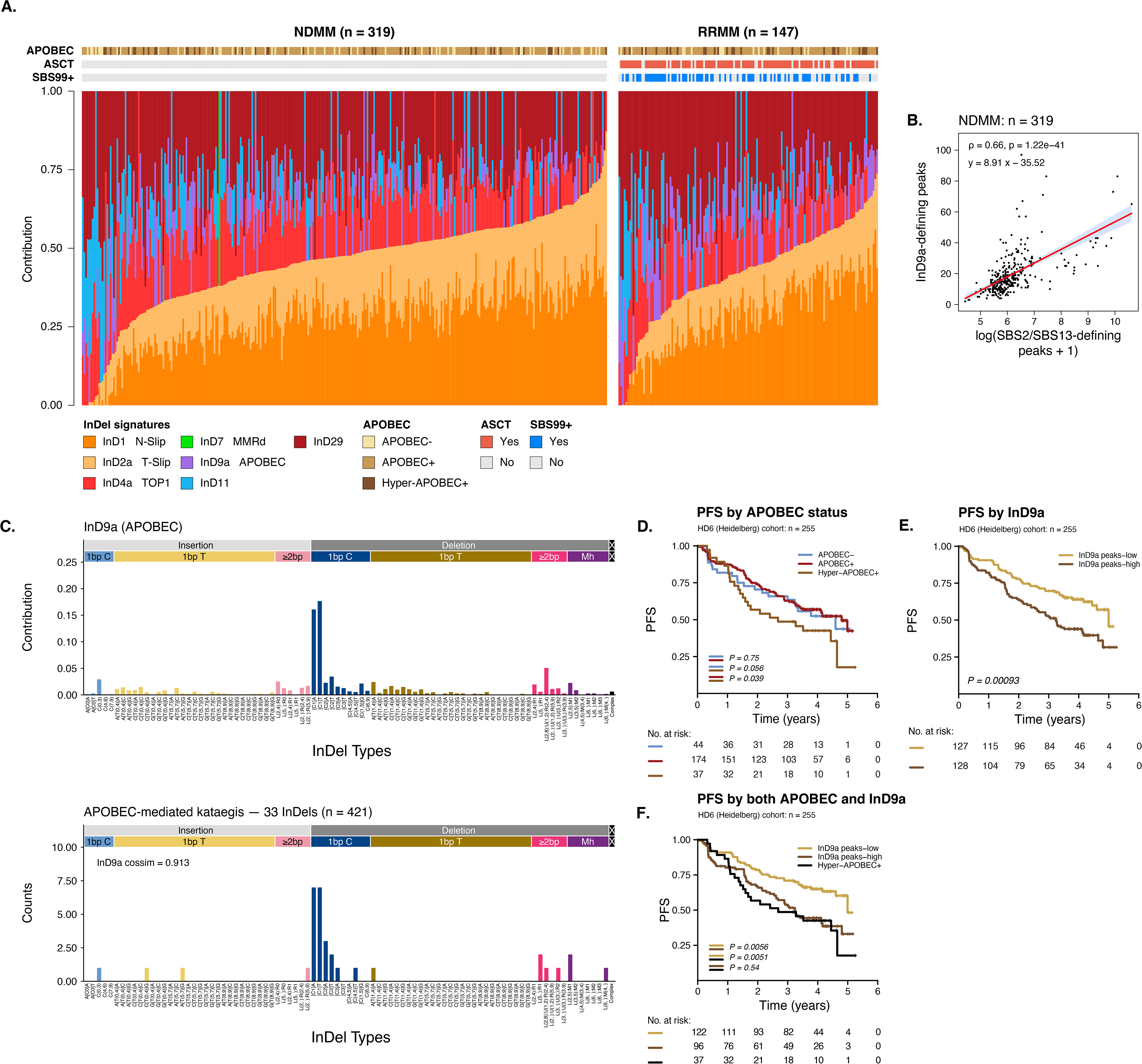
InD9a captures APOBEC-associated mutagenesis and improves prognostic stratification in MM. **(A)** Relative contributions, after *mmsig* fitting of the seven reference InDel mutational signatures across the discovery cohort (n = 466), stratified by NDMM (n = 319) and RRMM (n = 147). Each bar represents an individual sample, with colors indicating the relative contribution of each signature. Samples are ordered by increasing sum of InD1 and InD2a contribution within each disease setting. Annotations indicate sample collection timepoint, APOBEC status based on SBS2/SBS13 contributions, ASCT status, and the presence of SBS99, associated with melphalan exposure. **(B)** Correlation between the number of InD9a-defining InDel peaks and SBS2/SBS13-defining peaks (**Supplementary methods**) in NDMM samples (n = 319). Each point represents an individual sample. The fitted regression line and its confidence interval are shown, together with the Spearman correlation coefficient and associated p value. Values are log-transformed as log10(count + 1). **(C)** Comparison of the 89-channel mutational profile of the reference InD9a signature (top) and the distribution of 33 InDels identified within 293 APOBEC-mediated kataegis events (bottom). The cossim between the two profiles is indicated (0.913). **(D–F)** Kaplan–Meier estimates PFS in the Heidelberg GMMG-HD6 cohort (n = 255), stratified by SBS-based APOBEC status (**D**), InD9a burden (**E**), and a combined classification incorporating SBS-based hyper-APOBEC status and InD9a burden (**F**). InD9a burden was dichotomized using the cohort median. P values are displayed in each panel, with numbers at risk provided below the survival curves. SBS: single-base substitution; ASCT: autologous stem-cell transplantation; PFS: progression-free survival.

We posited that isolating the characteristic mutational peaks of these signatures (**InD9a:** [Del(C):R1]T and [Del(C):R1]A, **SBS2:** T[C>T]A and T[C>T]T, **SBS13:** T[C>G]A and T[C>G]T; **Supplementary Table 9; Supplementary Methods**) might decrease background noise and, indeed, the numbers of InD9a-defining and SBS2/SBS13-defining mutations exhibited a substantially stronger correlation (Spearman’s ρ=0.66, p<0.001; **Figure 3B**). This may suggest a degree of background activity or signature bleeding in the APOBEC indel signature that may dilute the specific mutational processes. We further investigated the association between InD9a and APOBEC by examining InDels occurring within regions of APOBEC-associated kataegis, as previously performed^46^. Briefly, kataegis events represent clusters of hypermutated regions characterized by at least six somatic SNVs with an average intermutational distance (IMD) of less than 1,000 bp^17^. Among 421 samples (1 sample per patient) in the discovery cohort, we identified 293 APOBEC-mediated kataegis events harboring in total 33 InDels, whose mutational profile exhibited strong similarity to InD9a (cossim = 0.913; **Figure 3C**, **Supplementary Figure 5, Supplementary Methods**), providing additional evidence to link InD9a to APOBEC activity.

Given the established association of APOBEC mutagenesis with adverse clinical outcomes in MM, including inferior progression-free survival (PFS)^6,20,21^ we next investigated the clinical relevance of InD9a in patients treated homogeneously in the GMMG-HD6 cohort (n = 255)^34,35^. Previous studies have demonstrated that patients with hyper-APOBEC activity (SBS2+SBS13 > 11%) experience inferior PFS^6,11,43,47^. However, below this threshold, presence or absence of detectable APOBEC mutagenesis is not informative for clinical outcome (**Figure 3D**). However, we observed that patients harboring higher InD9a burden (above the median) experienced inferior PFS (Log-rank test, p < 0.001; **Figure 3E**). To investigate whether InD9a could provide additional prognostic information beyond conventional SBS-based classification, we used InD9a-defining mutation burden to stratify non-hyper-APOBEC patients (Log-rank test, p = 0.0012; **Figure 3F**). Notably, patients with high InD9a burden who did not meet the SBS-based hyper-APOBEC definition (≤11%) experienced similarly unfavorable outcomes to those classified as hyper-APOBEC, whereas patients with low InD9a burden exhibited more favorable PFS. Together, these findings suggest that APOBEC-associated InDel mutagenesis captures clinically relevant activity not fully reflected by SBS-based classification and may improve prognostic stratification in MM.

### InD29 is associated with structural variant breakpoints and enriched in multiple myeloma with melphalan-induced mutagenesis

We next investigated InD29, a signature sharing substantial similarities with InD8, which has been associated with NHEJ-mediated repair of DNA double-strand breaks (DSBs). We previously identified the NHEJ-linked COSMIC ID8 signature exclusively in relapsed lymphoid malignancies, particularly following exposure to radiotherapy or mutagenic chemotherapy, including melphalan and platinum therapy (**Supplementary Figure 6**)^32^. Here, however, we detected the analogous InD29 as active in both NDMM and RRMM, reflective of the increased resolution of the new InDel classification schema to identify more subtle mutational signature contributions. Nevertheless, as expected, InD29 contributions were significantly higher in RRMM compared to NDMM (Wilcoxon p < 0.001; **Figure 4A**), as well as in samples harboring the melphalan-mediated SBS99 signature (Wilcoxon p < 0.001; **Figure 4B**). Given the proposed association between InD29 and repair of DSB/NHEJ, we next investigated its relationship with SVs. Patients above the median burden of InD29-defining InDel classes (**InD29:** A[Del(T):R(1,4)]A, Del(5,):R1 and Del(6,):M1; **Supplementary Methods**) harbored significantly more SVs (Wilcoxon p = 5.27 × 10^-3^; **Figure 4C**). Furthermore, controlling for patients with focal complex SV with multiple breakpoints (such as chromoplexy and chromothripsis)^27,48,49^, InD29-enriched cases exhibited a broad genomic distribution of SV breakpoints, as reflected by a greater number of 1-Mb genomic bins containing at least one SV breakpoint (p = 1.08 × 10^-2^; **Figure 4D**). Together, these findings support an association between InD29-related mutagenesis and [failure to repair] SVs.

**Figure 4.**
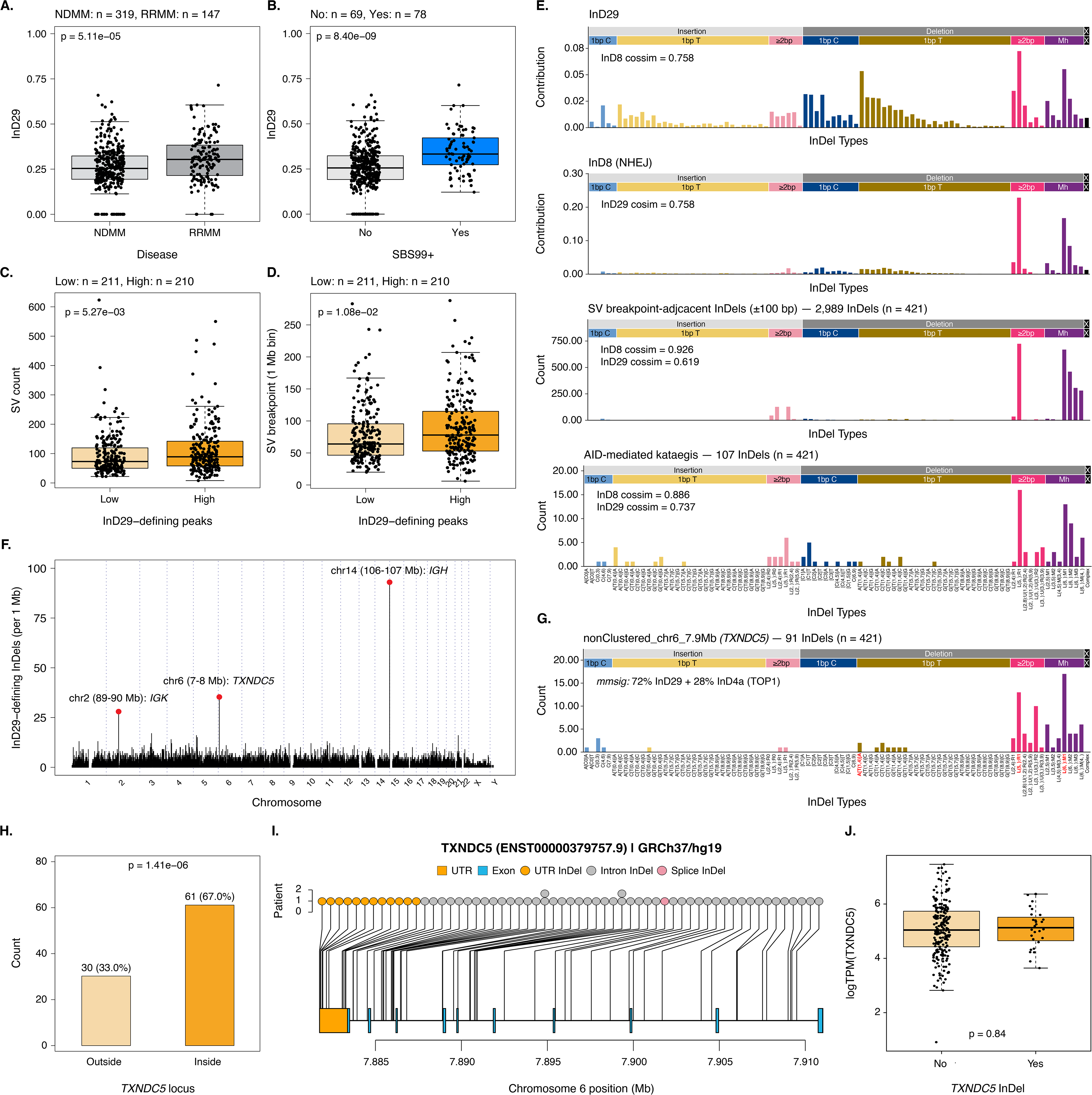
InD29 is associated with SVs and identifies recurrent InDel hotspots in MM. **(A)** Comparison of InD29 signature contributions between NDMM (n = 319) and RRMM (n = 147) samples in the discovery cohort. **(B)** Comparison of InD29 contributions between samples with (n = 78) and without (n = 69) the melphalan-associated signature SBS99 in RRMM. **(C)** Comparison of total SV burden and **(D)** the number of SV breakpoints across non-overlapping 1-Mb genomic bins between samples with low (n = 211) and high (n = 210) InD29-defining InDel burdens. Samples were stratified according to the median InD29-defining InDel burden. **(E)** Comparison of the 89-channel mutational profiles of the reference InD29 signature, the NHEJ-associated InD8 signature, InDels located near SV breakpoints (±100 bp), and InDels identified within AID-mediated kataegis events in the discovery cohort (n = 421). Cossim between InD8 and InD29 reference signatures are indicated. **(F)** Genome-wide distribution of InD29-defining InDels across non-overlapping 1-Mb genomic bins, highlighting recurrent enrichment at the immunoglobulin loci *IGH* and *IGK* as well as the *TXNDC5* locus on chromosome 6. **(G)** Distribution of 91 somatic InDels identified within the *TXNDC5* genomic region in the discovery cohort (n = 421). Signature fitting via *mmsig* identified contributions from InD29 (72%) and InD4a (28%). **(H)** Distribution of InDels located within (n = 61) or outside (n = 30) the *TXNDC5* locus within the chromosome 6 hotspot region. **(I)** Genomic distribution and functional annotation of InDels affecting *TXNDC5* (ENST00000379757.9; GRCh37/hg19). Individual InDels are mapped to their genomic positions, with the number of affected patients indicated. **(J)** Comparison of *TXNDC5* expression within the GMMG-HD6 cohort, measured as logTPM, between samples with and without *TXNDC5* InDels. Each point represents an individual sample. For boxplots, boxes indicate the interquartile range, with the central line representing the median. P values are displayed in the corresponding panels. SV: structural variant; Mb: megabase; AID: activation-induced cytidine deaminase; UTR: untranslated region; TPM: transcripts per million.

To further investigate the mutational processes underlying InD29, we compared its 89-channel InDel profile with that of InD8, confirming substantial similarity between the two signatures (cossim = 0.758; **Figure 4E**). Given the association with SV burden, we subsequently examined the InDel spectrum of mutations occurring within ±100 bp of SV breakpoints (**Supplementary Figure 7; Supplementary Methods**). Across 421 patients, we identified 2,989 InDels near SV breakpoints, whose mutational profile exhibited greater similarity to InD8 (cossim = 0.926) than to InD29 (cossim = 0.619; **Figure 4E**), suggesting InD29 may be composed of more than one mutational process. We next examined InDels occurring within regions of AID-mediated kataegis, another established source of localized somatic hypermutation in MM^9^. The mutational spectrum of 517 AID-mediated kataegis events harboring 107 InDels resembled InD8 (cossim = 0.886), while also exhibiting similarity to InD29 (cossim = 0.737; **Figure 4E**; **Supplementary Figure 5; Supplementary Methods**). These findings indicate that AID mutagenesis generates DNA DSB that NHEJ attempts to repair, generating a mutational footprint similar to what is observed with ionizing radiation and some mutagenic chemotherapies.

### TXNDC5 is an InDel hotspot enriched for TOP1 and NHEJ-mediated mutagenesis

We next investigated the genome-wide distribution of InD29-defining InDels to identify regions of positive selection. By quantifying these InDels across non-overlapping 1-Mb genomic bins, we identified prominent peaks at the immunoglobulin loci IGH and IGK (**Figure 4F**, **Supplementary Table 10**). The next most prominent peak was located at chromosome 6p24.3, encompassing the *TXNDC5* locus, with a total of 30 InD29-defining InDels. We then applied a piecewise constant fitting (PCF) approach to orthogonally validate the *TXNDC5* region as an InDel hotspot (**Supplementary Table 11**)^27^. *TXNDC5* is highly expressed in MM and resides within a super-enhancer-associated genomic region at chromosome 6p24.3 that has previously been identified as a recurrent partner of *MYC* rearrangements and as an SV hotspot^27,50,51^. The InDel spectrum at the *TXNDC5* locus was reconstructed predominantly by InD29 (72%), with a smaller contribution from the TOP1-associated signature InD4a (28%; **Figure 4G**). These findings may reflect both the instability and the high transcriptional activity at *TXNDC5* with replication-associated mutagenesis (**Supplementary Figure 8**)^52^.

Of the 91 InDels identified within the hotspot, 61 (67.0%) mapped within the *TXNDC5* gene itself, whereas the remaining 30 (33.0%) occurred in the surrounding genomic region (Fisher’s p < 0.001; **Figure 4H**). Intragenic InDels were predominantly located within introns and the 3′ untranslated region (3′UTR)(**Figure 4I**, **Supplementary Table 12**). To explore the potential functional consequences of these alterations, we examined available RNA-sequencing (RNA-seq) data from NDMM patients enrolled in the GMMG-HD6 trial, but no significant difference in *TXNDC5* expression was observed between samples harboring *TXNDC5* InDels and those without such events (Wilcoxon p = 0.84; **Figure 4J**). Nevertheless, given the accumulation of InDels within the 3′UTR, we used TargetScan to investigate predicted microRNA (miRNA) binding sites within this region^53^. This analysis identified predicted binding sites for miR-129-5p and other miRNAs, including members of the miR-320 family, some of which overlapped the observed InDels (**Supplementary Table 13**). Although these alterations were not associated with detectable changes in overall *TXNDC5* expression, their localization raises the possibility that a subset may disrupt miRNA-mediated post-transcriptional regulation.

### Independent validation of InDel mutational signatures

To independently validate InDel mutational processes active in MM, we applied the same *de novo* signature extraction and fitting framework to the MIDAS cohort (n = 421)^36,37^. Five *de novo* mutational signatures were extracted and delineated into seven reference signatures (**Figure 5A**, **Supplementary Figure 9-10, Supplementary Tables 14-18**). The extracted signatures recapitulated the principal mutational processes observed in the discovery cohort, including replication slippage, TOP1-mediated mutagenesis, APOBEC, and NHEJ. Notably, InD2b was identified in place of InD2a, previously described as a variant of InD2a^33^. InD11 (of unknown biological significance) was the only signature not observed, whereas an additional signature associated with HRD (InD6) was identified. The identification of InD6 was consistent with previous genomic characterization of the MIDAS cohort, which identified a rare HRD *BRCA2*-mutant case^37^. We were able to recapitulate the association of InD29 InDels and SVs within the validation cohort, as well as reproduce the relative concordance of InD9a with SBS2/SBS13 activity, further supporting InDels ability to capture a complementary component of APOBEC mutagenesis (**Supplementary Figures 11-12**). As the achievement of measurable residual disease (MRD)-negativity (10^-6^) prior to maintenance therapy was the primary endpoint of the MIDAS trial^36^, we specifically evaluated the relationship between InD9a contribution and MRD status. Notably, patients who remained MRD-positive following induction therapy at 10^-5^ exhibited significantly higher InD9a compared with those achieving MRD negativity (Wilcoxon p < 0.001; **Figure 5B**). This association remained when MRD status was evaluated at the more stringent threshold of 10^-6^ (Wilcoxon p < 0.001; **Figure 5C**). Together, these findings establish the reproducibility of the major InDel mutational processes and their biological associations across independent MM cohorts, while identifying InD9a as a poor prognostic feature across various treatment paradigms.

**Figure 5.**
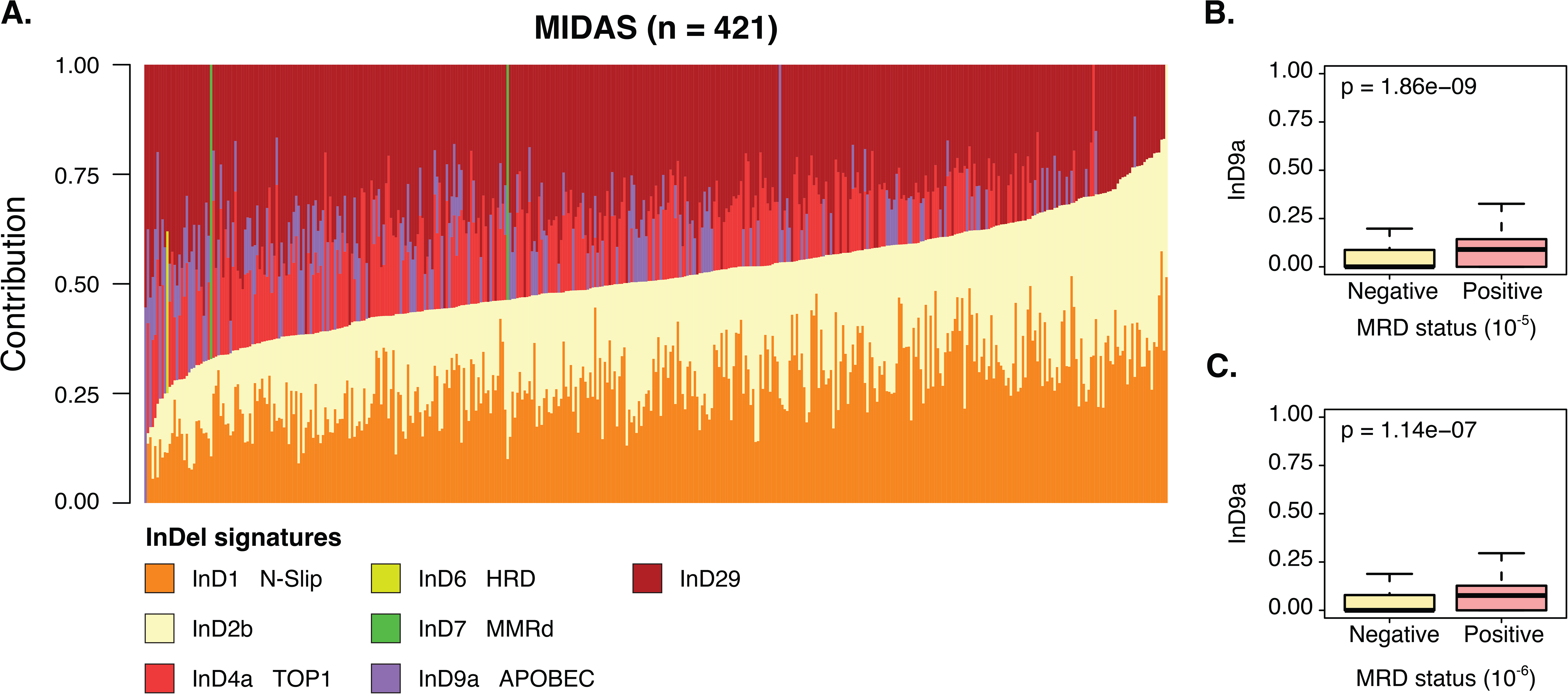
Validation of InDel mutational signatures and association of InD9a with MRD status in the MIDAS cohort. **(A)** Relative contributions, after *mmsig* fitting of the seven reference InDel mutational signatures across the validation cohort (n = 421), consisting of NDMM samples. Each bar represents an individual sample, with colors indicating the relative contribution of each signature. Samples are ordered by increasing sum of InD1 and InD2b contribution within each disease setting. **(B–C)** Comparison of InD9a signature contributions according to post-induction MRD status, assessed at sensitivity thresholds of 10 (**B**) and 10 (**C**). Boxes indicate the interquartile range, with the central line representing the median. P values are displayed in each panel. HRD: homologous recombination deficiency; MMRd: mismatch repair deficiency; MRD: minimal residual disease.

### Identification of rare DNA repair-deficient phenotypes in MM

Within the discovery cohort, we identified one sample (MM165_BL) exhibiting a markedly elevated InDel burden (n = 8,257), for which InD7 accounted for 62.0% (**Figure 3A**; **Figure 6A**, **top panel; Methods**). This sample was also characterized by an elevated SNV burden (n = 14,306), but notably lacked detectable contributions from SBS2 and SBS13, which are frequently associated with hypermutated phenotypes in MM. Instead, SBS signature analysis revealed a substantial contribution from the MMRd signature SBS6 (∼25%; **Supplementary Figure 13**). Genomic characterization revealed a dominant pattern of aneuploidy consisting of extensive chromosomal deletions, including that of *MSH2* and *MSH6* on chromosome 2, and a low estimated ploidy of 1.57 (i.e., hyperhaploid; **Figure 6A**; **bottom panel**)^54,55^. Notably, no pathogenic germline variants were identified in either gene, suggesting that the MMRd may have arisen through another mechanism.

**Figure 6.**
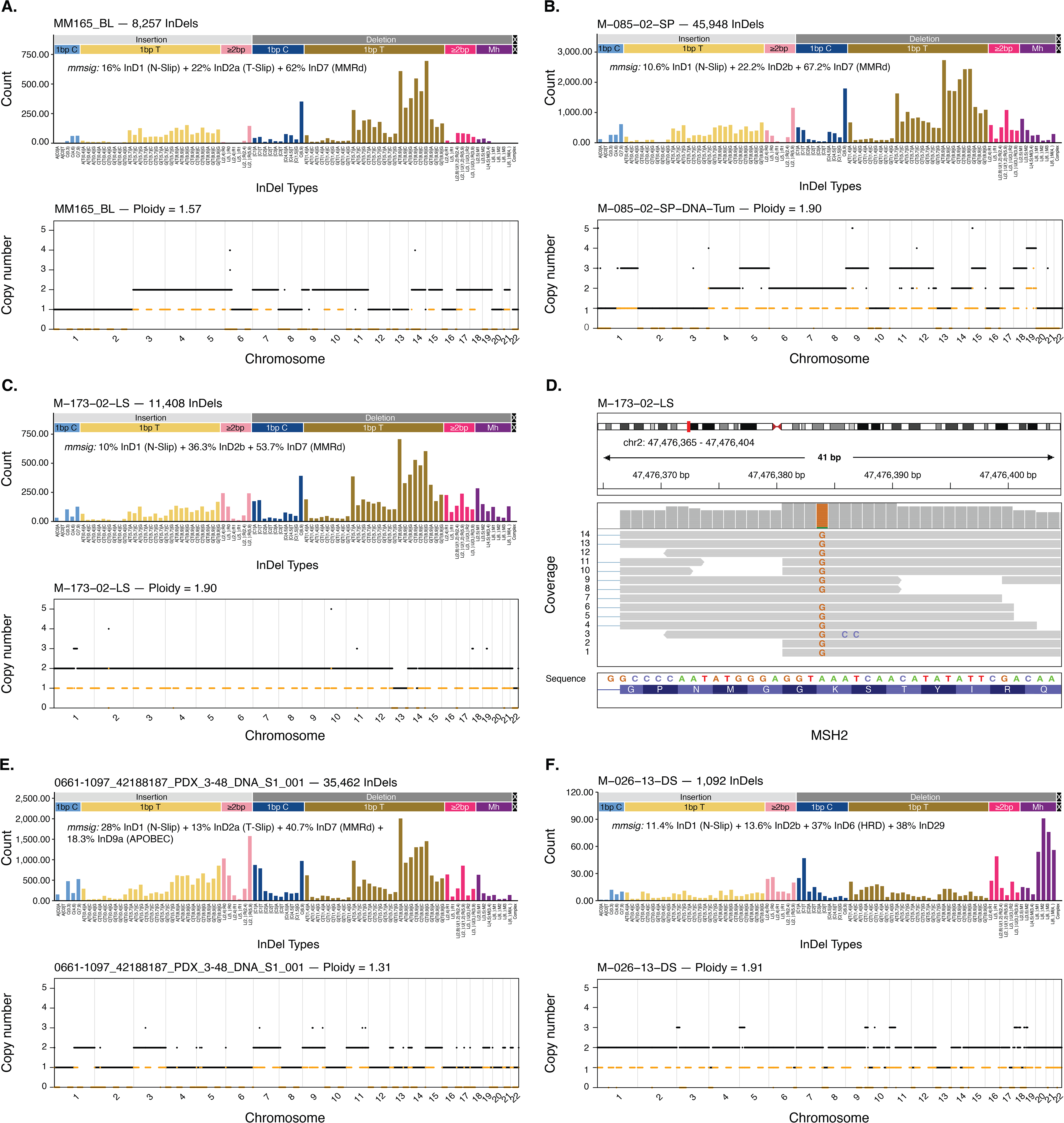
InDel mutational signatures identify rare DNA repair-deficient phenotypes in MM. **(A–C)** Somatic InDel mutational profiles (top) and genome-wide copy-number profiles (bottom) of three MM samples exhibiting MMRd-associated mutational signatures, including one sample from the discovery cohort (MM165_BL; **A**) and two samples from the independent MIDAS validation cohort (M-085-02-SP; **B**, and M-173-02-LS; **C**). InDel profiles are displayed according to the 89-channel classification framework, with bar heights representing the number of InDels assigned to each mutational channel. The total InDel burden and relative contributions of fitted reference signatures via *mmsig* are indicated for each sample. CNV profiles illustrate the distribution of chromosomal gains and losses, with estimated tumor ploidy indicated. The solid black lines represent total copy number and the yellow dashed lines represent minor copy number. **(D)** IGV visualization of RNA-seq reads aligned to the *MSH2* locus in sample M-173-02-LS, highlighting a sequence alteration affecting *MSH2*. **(E)** Somatic InDel profile (top) and CNV profile (bottom) of a hyperhaploid PDX model (0661-1097_42188187_PDX_3-48_DNA_S1_001). InDel signature fitting was performed using all extracted signatures from both validation and discovery cohort. **(F)** Somatic InDel mutational profile (top) and CNV profile (bottom) of an HRD-associated sample from the MIDAS validation cohort (M-026-13-DS). CNV: copy number variation; IGV: Integrative Genomics Viewer; RNA-seq: RNA sequencing; PDX: patient-derived xenograft.

Within the MIDAS cohort, two additional samples previously identified as MMRD were dominated by InD7. The presence of MMRd was also confirmed via PRRDetect (**Methods**). The first sample (M-085-02-SP) harbored 45,948 InDels (**Figure 5A**, **Figure 6B**; **top panel**). This sample carried a germline *MSH2* Q339* mutation accompanied by copy-neutral loss of heterozygosity (LOH) and likewise displayed a hyperhaploid profile (**Figure 6B**; **bottom panel**). The second sample (M-173-02-LS) harbored a somatic *MSH2* K675E mutation, previously described as potentially pathogenic, with a prominent InD7 contribution (53.7%; **Figure 5A**, **Figure 6C**). While the WT allele appeared intact, examination of sequenced RNA revealed most informative reads supported the somatic *MSH2* K675E, with no expression of the WT allele (**Figure 6D**), suggesting epigenetic silencing of the WT allele. Further supporting this possibility, we examined a recently characterized patient-derived xenograft (PDX) model of a MM exhibiting a MMRd phenotype^56^. This case displayed a dominant InD7 InDel landscape, and hyperhaploid CNV profile that was characterized by monoallelic deletion of *MSH2* (**Figure 6E**). Using methylation data derived from long-read WGS data (PacBio) we identified that the monoallelic loss of *MSH2* was accompanied by epigenetic silencing of the remaining allele through promoter hypermethylation and gene-body hypomethylation resulting in loss of *MSH2* expression and low protein abundance (**Supplementary Figure 14**). As observed in other cancers, we identify *MSH2* promoter hypermethylation to be associated with loss of MSH2 expression in MM^57,58^. Overall, MMRd MM is a recurrent phenotype achieved via a variety of genetic and epigenetic mechanisms and is associated with hyperhaploidy.

Finally, we investigated the HRD-associated mutational signature identified during *de novo* extraction in the MIDAS cohort. As previously mentioned, sample M-026-13-DS carried a germline frameshift mutation in *BRCA2* (T1858fs). This was accompanied by deletion of the WT allele and was previously seen to be enriched for focal deletions^37^. Harboring 1,092 InDels, InD6 accounted for 37.0% of its signature contribution (**Figure 5A**, **Figure 6F**; **Supplementary Methods**). In conclusion, these findings suggest that InDel mutational signatures assist in identification of rare and underrecognized subsets of MM characterized by underlying DNA damage repair deficiencies, even in the absence of clear germline variants, and identifies potential therapeutic vulnerabilities.

## DISCUSSION

While the SBS landscape of MM has been extensively analyzed^9,20,21,27,39^, the process shaping the accumulation of InDels have been previously underexplored. In this study, we comprehensively characterized the landscape of somatic InDels in MM using WGS data from 887 samples. The identification of consistent mutational processes across these clinically heterogeneous cohorts, including an independent validation cohort, supports the robustness and generalizability of our findings. By implementing a recently developed 89-channel InDel classification schema^33^, we identified distinct mutational processes associated with replication-mediated slippage, APOBEC activity, DSB repair and NHEJ, as well as rare DNA damage repair deficiencies, that shape the mutational landscape of the disease. Importantly, our findings demonstrate that InDel mutational signatures provide complementary biological and clinical information beyond conventional SBS-based analyses, while revealing previously unrecognized relationships between InDels and structural rearrangements.

One of the most clinically relevant findings is the identification of APOBEC-associated InDel mutagenesis through InD9a. Although InD9a correlated with SBS2/SBS13, considerable discordance between their relative contributions suggests that SBS-based classification does not fully capture the spectrum of APOBEC-associated mutagenesis, particularly single-base cytosine deletions occurring in APOBEC-associated sequence contexts. Notably, InD9a burden identified patients with inferior outcomes who otherwise would remain indistinguishable using conventional SBS-based APOBEC assignment. These findings suggest that integrating InDel and SBS signatures may provide a more comprehensive assessment of APOBEC activity and improve prognostic classification as established across cohorts in two randomized clinical trials^34,36^. The increased resolution provided by the 89-channel classifier allowed for another important observation in the identification of InD29 and NHEJ-mediated mutagenesis in both NDMM and RRMM. Our previous work identified this exclusively in RRMM following exposure to radiotherapy or mutagenic chemotherapy, because an extreme contribution was required for detection using the previous COSMIC 83 channel classifier^15,32^. We established InD29’s association with SV burden and genomic breakpoint dispersion, supporting a link with NHEJ-mediated repair of DSB formation and repair ^41^. It follows that this process would therefore be amplified following accumulation of DSB due to mutagenic therapy but demonstrates this as an endogenous process associated with continuous accumulation of structural variation. Furthermore, the enrichment of InD8/InD29-like mutations within AID-associated kataegis regions raises the possibility that aberrant AID activity contributes to these mutational patterns^9^.

The identification of *TXNDC5* as a recurrent InDel hotspot provides additional evidence linking InDel mutagenesis to regions of genomic instability and/or high transcriptional activity. Although this locus has previously been implicated in complex rearrangements^18,50^, the enrichment of InDels, particularly those associated with InD4a and InD29, suggests that multiple mutational processes may contribute to its genomic instability. Given the high transcriptional activity of *TXNDC5* and its potential relevance to plasma cell biology, these findings raise the possibility that this locus represents a biologically important vulnerability in MM warranting further investigation. Finally, the identification of MMRd- and HRD-associated InDel signatures highlights their utility in recognizing rare DNA repair-deficient phenotypes in MM. The detection of these signatures in independent cohorts, including MMRd-associated profiles without identifiable pathogenic germline alterations, suggests that InDel analysis may reveal functional repair deficiencies not readily apparent from conventional genomic characterization, and may identify patients for whom personalized therapy, such as with checkpoint blockade, might be revisited^59–61^. Furthermore, identification of the MMRD phenotype as being enriched in hyperhaploid myeloma suggests this poor-risk subgroup is worthy of further interrogation for this purpose^54,55^.

In summary, our study provides a comprehensive characterization of the InDel mutational landscape in MM, demonstrating that these mutations capture previously underappreciated endogenous biological processes and patterns of genomic instability involved in shaping the evolution of the disease. Although the precise mechanisms underlying some of these signatures remain incompletely understood, our findings highlight the importance of incorporating InDel mutagenesis into future studies of MM evolution. Further investigations integrating genomic, transcriptomic, and functional approaches will be essential to elucidate the biological origins of these mutational processes and refine prognostic stratification of MM.

## SOURCE CODE

The source code for the modified *mmsig* framework specifically for InDel signature fitting is publicly available at https://github.com/UM-Myeloma-Genomics/indel_fitting.

## DATA AVAILABILITY

Whole-genome sequencing (WGS) data from the GMMG-HD6 trial (NCT02495922) are available through the European Genome-phenome Archive (EGA) under accession number EGAS00001007469 (datasets EGAD50000000681, EGAD50000000682, and EGAD50000000683). Additional WGS data from patients with NDMM and RRMM treated at Heidelberg University are available under accession numbers EGAS00001006538, EGAS00001004363, EGAS00001004805, and EGAS00001005973.

WGS data from the independent IFM-MIDAS validation cohort are available through EGA under accession number EGAS50000001573 (datasets EGAD50000002251, EGAD50000002687, and EGAD50000002807).

Data from SCCC and IMR was newly sequenced and will be uploaded upon acceptance of the manuscript for publication.

## ACKNOWLEDGMENTS

A.M.P. is funded by the Medical Data Scientist Program of Heidelberg University, Faculty of Medicine. NW is funded by the Advanced Medical Scientist Program of Heidelberg University, Faculty of Medicine.

Ola Landgren is supported by Paul J. DiMare Endowed Chair in Immunotherapy, Paula and Rodger Riney Foundation, Tow Foundation, Myeloma Solutions Fund, Rising Tide Foundation, Blood Cancer United (formerly: Leukemia & Lymphoma Society), and Cannon Guzy Family Fund.

B.A.W. is funded by Blood Cancer United (grant 8049-25) and the U.S. Department of Defense (CA230482).

Benjamin Diamond is supported by the American Cancer Society Clinician Scientist Development Grant under Award Number CSDG-26-1595952-01-CDP-W25.

We thank all patients and families for participating and contributing to the study.

The Heidelberg Team thanks the Sample Processing Lab (SPL), the High Throughput Sequencing unit of the Genomics & Proteomics Core Facility and the Omics IT and Data Management Core Facility (ODCF) of the German Cancer Research Center (DKFZ), the DKFZ-Heidelberg Center for Personalized Oncology (DKFZ-HIPO) office, the Biobank Multiple Myeloma UKHD and the Myeloma Registry for excellent services. Support and funding of the project via the Dietmar-Hopp Foundation and the NCT Heidelberg Molecular Precision Oncology Program (project H067 and K08K) is gratefully acknowledged. Data storage service via SDS@hd is supported by the Ministry of Science, Research and the Arts Baden-Württemberg (MWK) and the German Research Foundation (DFG) through grants INST 35/1314-1 FUGG and INST 35/1503-1 FUGG.

Research reported in this publication was performed in part at the Onco-Genomics Shared Resource (SCR_022502), Cancer Modeling Shared Resource (SCR_022891), Biospecimen Shared Resource (SCR_022889), and Cytometry and Imaging Shared Resource (SCR_022501) of the Sylvester Comprehensive Cancer Center at the University of Miami, which is supported by the National Cancer Institute (NCI) of the National Institutes of Health (NIH) under award number P30CA240139.

