## Supplementary Methods for "A refined InDel taxonomy broadens the repertoire of mutational processes shaping multiple myeloma"

*Sample collection, and library preparation*

For the IMR cohort, BM aspirates underwent red blood cell lysis and mononuclear cell isolation using Ficoll density gradient centrifugation, followed by CD138 magnetic bead enrichment (Miltenyi Biotec) of tumor plasma cells. Matched germline controls were obtained from peripheral blood or saliva. Tumor DNA was extracted using the AllPrep DNA/RNA/miRNA Universal Kit (Qiagen), while germline DNA was isolated using magnetic bead-based purification (Chemagic 360, PerkinElmer). WGS was performed using Illumina short-read sequencing, achieving an average coverage of 65.5× for tumor samples and 33.2× for matched germline controls.

For the SCCC cohort, BM samples with sufficient plasma cell infiltration, as determined by clinical flow cytometry, underwent tumor plasma cell enrichment using CD138 magnetic microbeads or fluorescence-activated cell sorting. DNA was extracted using automated magnetic bead-based purification (KingFisher Apex, Thermo Fisher Scientific). WGS libraries were prepared using the Roche KAPA EvoPlus V2 kit, following PCR-free or PCR-based workflows. Samples with insufficient DNA input underwent whole-genome amplification using the BioSkryb ResolveDNA V2 system prior to library preparation. Libraries were sequenced on the Illumina NovaSeq X Plus platform using 150-bp paired-end reads, achieving an average coverage of 80× for tumor samples and 40× for matched germline controls. Sequencing data were demultiplexed using Illumina BCL Convert.

For the independent validation cohort, we included WGS data from 421 patients with NDMM enrolled in the MIDAS trial (IFM) (1). Sample collection, tumor plasma cell enrichment, DNA extraction, library preparation, and sequencing procedures were performed as previously described by Magrangeas et al., 2026 (2).

*Next generation sequencing pipeline*

WGS data were processed using different bioinformatic pipelines, reference genome assemblies, and somatic calling procedures, as detailed below. For the Heidelberg University cohort, raw sequencing reads were processed and aligned to the human reference genome GRCh37 (hg19; hs37d5) using the DKFZ OTP WGS pipeline. Somatic SNVs were identified using SAMtools mpileup (v1.2.166-3) (3), while InDels were called using Platypus (4). Additional filtering was applied to both variant classes, including the removal of variants overlapping blacklisted genomic regions and immunoglobulin loci, with further SNV-specific filtering performed using fpfilter (5). CNV were identified using ASCAT (6), and SVs were called using SOPHIA. For the RNA-seq data, sequencing reads were aligned to GRCh37 (hg19; hs37d5) using STAR (v2.5.2b) (7), and gene-level counts were quantified using featureCounts (Subread v1.5.1) (8).

For both the IMR cohort and the SCCC cohort, all samples were aligned to the human reference genome GRCh38 (hg38) and analyzed using the MGP1000 pipeline (https://github.com/UM-Myeloma-Genomics/MPG). SNVs were identified by MuTect2, VarScan2, and Strelka, while InDels were identified by VarScan2, MuTect2, SvABA, and Strelka (at least 3 out of 4 callers). CNVs were evaluated using ASCAT. SVs were defined by Manta, DELLY, and SvABA and IgCaller and passed through additional quality filters as previously described.

For the MIDAS IFM validation cohort, WGS reads were aligned to the human reference genome GRCh38 (hg38), and SNVs, CNVs, and SVs were identified as previously described (2). Somatic InDels were called using Illumina DRAGEN (v4.3.6) (9), retaining variants classified as PASS or weak_evidence with a somatic quality score (SQ) ≥10, followed by additional filtering based on sequencing depth, variant allele frequency (VAF), and tumor–normal evidence. Finally, as an additional dataset for cross-tumor validation of InDel mutational signatures, we included 113 samples from BRCA-EU breast cancer cohort, included in the Pan-Cancer Analysis of Whole Genomes (PCAWG) project (https://dcc.icgc.org/) (10).

*InDel mutational signature extraction*

Somatic InDels were classified according to the 89-channel taxonomy developed by Koh et al. (2025) (11). InDels were assigned to their corresponding mutational channels to generate an 89-channel mutation count matrix for each sample. De novo InDel mutational signatures were extracted from the discovery cohort using non-negative matrix factorization (NMF). Signature extraction was performed using 20 bootstrap iterations, with 10 independent NMF repetitions per bootstrap, followed by hierarchical clustering of the extracted signatures. Solutions ranging from 1 to 10 signatures were evaluated, and the optimal solution was selected based on signature stability and reconstruction accuracy (**Supplementary Figures 3,9; Supplementary Tables 4-7,14-18**).

*Mutational signature fitting*

Mutational signatures were analyzed across all MM datasets. For SBS signatures, we used the R package *mmsig* (https://github.com/UM-Myeloma-Genomics/mmsig)(12). and fitted the established group of SBS signatures known to show activity in MM: SBS1 and SBS5 (Aging), SBS2 and SBS13 (APOBEC), SBS8 (Unknown), SBS9 [DNA polymerase eta (poly-eta) in germinal center (GC)], SBS18 [reactive oxygen species (ROS)] and SBS99 (SMS-MM1). SBS99 was fitted in RRMM samples exposed to melphalan, as previously employed (13). To identify hyper-APOBEC samples, we considered those with higher than 11% SBS2/SBS13 contribution (14).

To assign the de novo extracted signatures to established mutational processes, we performed pairwise fitting against the reference InDel signatures described by Koh et al., 2025 (11). For each de novo signature, all possible two-signature combinations were evaluated, and their relative contributions and cosine similarities (cossim) were calculated to identify the reference signatures providing the best reconstruction (https://github.com/UM-Myeloma-Genomics/Signature-Assignment). For each extracted signature, we selected the highest-ranking solution based on cossim reconstruction, with one exception. Within the discovery cohort for SIG_1, although the highest-ranking reconstruction involved InD1 (67%) and InD21 (33%; cossim = 0.969), we selected the second-best solution of InD1 (79%) and InD11 (21%; cossim = 0.966), as InD21 has been associated with concurrent POLD1 proofreading deficiency and mismatch repair deficiency (MMRd) (11). Given the absence of detectable *POLD1* mutations in our cohort, we considered the alternative reconstruction more biologically sound. Subsequently, reference InDel signatures were fitted to individual samples using a modified version of mmsig, adapted to accommodate the 89-channel InDel classification. Given the specific association of InD6 and InD7 with homologous recombination deficiency (HRD) and mismatch repair deficiency (MMRd), respectively, their fitting was restricted to samples exhibiting evidence of the corresponding deficiency. Specifically, InD6 was fitted exclusively to samples classified as HRD-positive by HRDetect (15), whereas InD7 was fitted only to samples classified as MMRd-positive by PRRDetect (11, 16). This was adopted based on our previous framework of SBS99 signature fitting in melphalan-exposed only samples.

*Mutational signature-defining peaks*

To identify the predominant mutational features (signature-defining peak) of each reference signature, we applied a Z-score–based approach to the normalized mutational probability profiles. For each signature, individual channel probabilities were standardized relative to the mean and standard deviation of all channels within that signature (96 channels for SBS signatures and 89 channels for InDel signatures). Mutational channels with a Z-score greater than 2 were considered predominant features of the corresponding signature. This approach identified two C>T channels in SBS2 (T[C>T]A and T[C>T]T), and two C>G channels in SBS13 (T[C>G]A and T[C>G]T). InD9a was defined by [Del(C):R1]T and [Del(C):R1]A, while InD29 was defined by A[Del(T):R(1,4)]A, Del(5,):R1, and Del(6,):M1 (**Supplementary Table 9**).

*InDel LiftOver*

For analyses requiring the aggregation of InDels across cohorts, such as kataegis analysis and InDel hotspot analysis via piecewise constant fitting (PCF; described below), genomic coordinates were converted from GRCh38/hg38 to GRCh37/hg19 using LiftOver to ensure consistency across datasets (17). GRCh37/hg19 was selected as the reference genome because it was used for the Heidelberg University (GMMG-HD6 trial) cohort, which constituted the largest proportion of the discovery cohort (n = 360/466).

*Kataegis analysis*

Kataegis events were identified using the katdetectr R package (18), defining kataegis as clusters of at least 6 SNVs with an average intermutational distance (IMD) of less than 1,000 bp. Mutational signature contributions within each event were estimated using an expectation-maximization (em) algorithm, considering the APOBEC-associated signatures SBS2 and SBS13 and the AID-associated signatures SBS84 and SBS85 (13). Individual kataegis events were subsequently classified as APOBEC-mediated or AID-mediated according to the predominant contribution of the corresponding mutational signatures (≥ 75% contribution; **Supplementary Figure 5**) (19).

*InDel proximity analysis to SV breakpoints*

To investigate the spatial relationship between InDels and SVs, we calculated the distance between each InDel and the nearest SV breakpoint. The distribution of InDel-to-SV breakpoint distances was examined across a 500 bp interval, revealing a pronounced accumulation of InDels immediately adjacent to SV breakpoints (**Supplementary Figure 7**). Based on these distributions, we selected a window of ±100 bp around SV breakpoints for identifying SV-adjacent InDels.

*InDel hotspot analysis*

To identify genomic regions exhibiting recurrent enrichment of somatic InDels, we adapted the PCF framework previously employed by Rustad et al., 2020 to identify SV hotspots in MM (20). Rather than analyzing SV breakpoint density, we applied this approach to the genomic distribution of somatic InDels to identify regions exhibiting localized accumulation of mutations relative to the surrounding genomic background. InDels were classified into eight categories based on the 89-channel InDel mutational signature framework: 1-bp C insertions, 1-bp T insertions, ≥2-bp insertions, 1-bp C deletions, 1-bp T deletions, ≥2-bp deletions, microhomology (Mh) deletions, and complex InDels.

*TXNDC5 InDel annotation*

InDels within the TXNDC5 locus were functionally annotated using ANNOVAR (21) with the Ensembl gene annotation database (hg19_ensGene) to determine their distribution across coding, intronic, splice-site, and untranslated (UTR) regions. To investigate the potential regulatory consequences of 3′-UTR InDels, predicted microRNA (miRNA)-binding sites were examined using TargetScan (**Supplementary Table 12**) (22).

*Data analysis and statistics*

Analysis was carried out in R version 4.2.3. Unless otherwise specified, we used the Wilcoxon rank-sum test to test for differences in continuous variables between two groups and Fisher’s exact test for 2 × 2 tables of categorical variables.
