## Supplementary Figures for "A refined InDel taxonomy broadens the repertoire of mutational processes shaping multiple myeloma"

**Supplementary Figure 1. Distribution of somatic InDels across the three cohorts.** Distribution of somatic InDels across the 89-channel classification framework in the Heidelberg (n = 360; 172,339 InDels), Indiana (n = 52; 21,212 InDels), and Miami (n = 54; 45,705 InDels) cohorts. Bar heights represent the aggregate number of InDels assigned to each mutational channel.

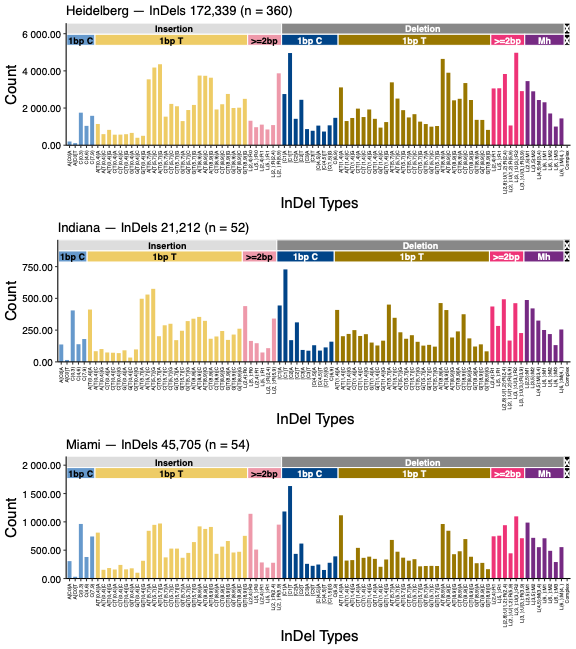

**Supplementary Figure 2. Relative contribution of 1-bp T deletions in NDMM and RRMM.** Comparison of the proportion of 1-bp T deletions relative to total InDel burden between NDMM (n = 319) and RRMM (n = 147) samples in the discovery cohort. Boxes indicate the interquartile range, with the central line representing the median. The P value is indicated.

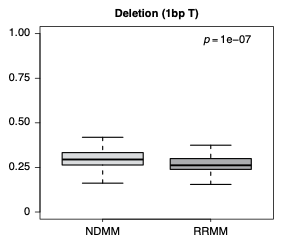

**Supplementary Figure 3. Evaluation of de novo InDel mutational signature extraction.** **(A)** Reconstruction accuracy and stability metrics across solutions containing 1–10 signatures using 20 bootstrap iterations for the discovery cohort (n = 466 samples). Lines represent normalized reconstruction error (solid black), reconstruction error against the original catalogue (dashed black), average silhouette width (red), minimum within-cluster cossim (green), maximum between-cluster cossim (blue), mean matched cossim (purple), and cophenetic correlation (orange). A four-signature solution was selected based on the balance between reconstruction accuracy and clustering stability. **(B)** Distribution of within-cluster cossim for the four-signature solution across three clustering methods: hierarchical clustering (hclust), matched clustering (MC), and partitioning around medoids (PAM). Boxplots illustrate the similarity of extracted signatures within each cluster, with median values indicated by horizontal black lines.

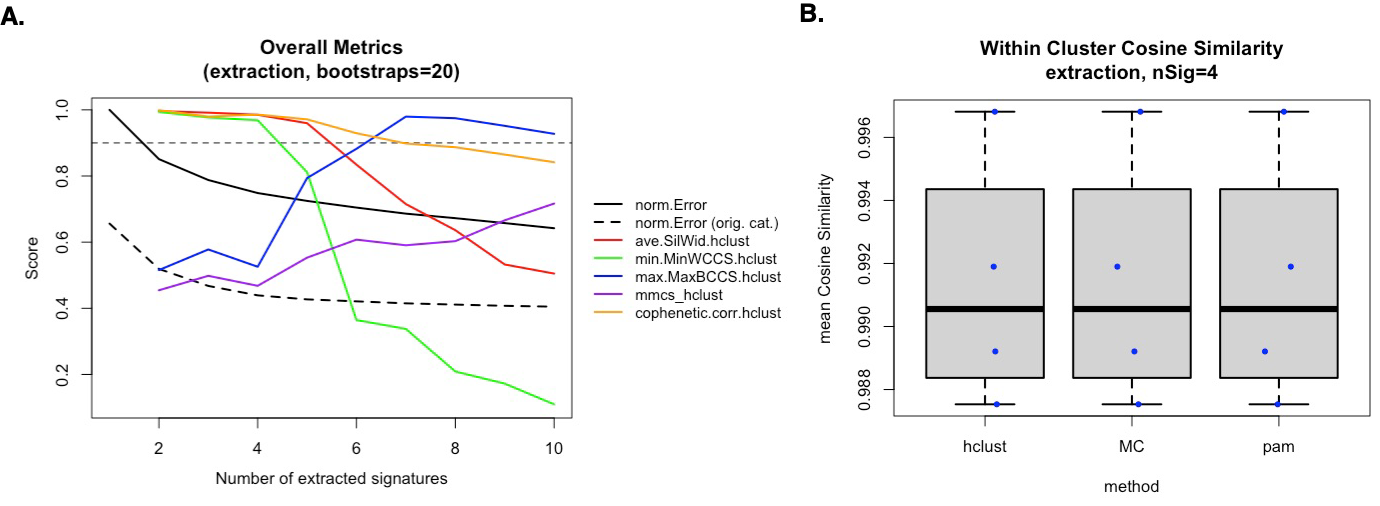

**Supplementary Figure 4. Association between InD9a and SBS2/SBS13.** **(A-B)** Scatterplots illustrating the relationship between InD9a and APOBEC-associated SBS2/SBS13 mutational activity, in **(A)** NDMM in the discovery cohort (n = 319) and **(B)** breast tumors in PCAWG (n = 113). Red lines indicate fitted linear regression models, and blue shaded regions represent 95% confidence intervals.

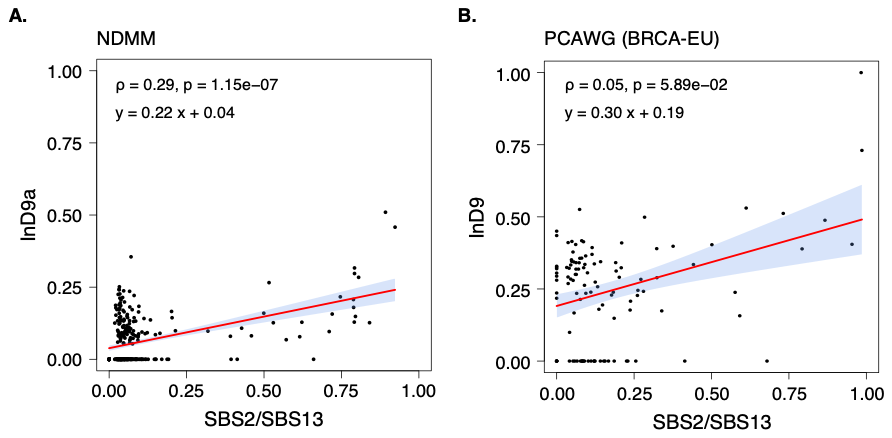

**Supplementary Figure 5. Genome-wide distribution of kataegis events in MM.** Stacked bar plot showing the distribution of kataegis events across the discovery cohort (421 samples), classified as APOBEC-mediated (red; n = 293), AID-mediated (blue; n = 517), or unassigned (gray; n = 81). The genome was divided into non-overlapping 1 Mb bins.

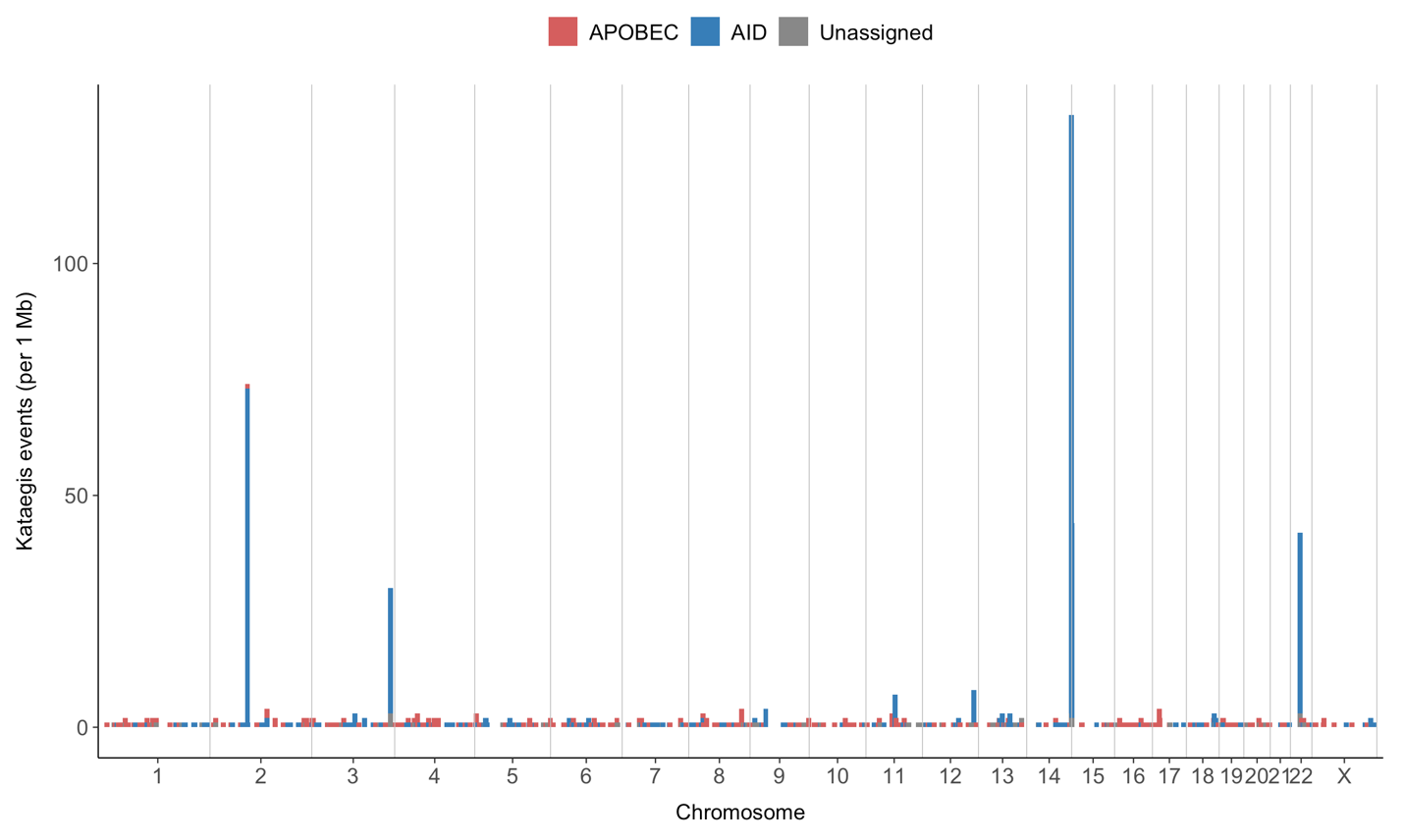

**Supplementary Figure 6. Correlation between InD29 and COSMIC ID8 mutational signature contributions in RRMM.** Scatter plot showing the relationship between the relative contributions of InD29 and the COSMIC ID8 mutational signature across RRMM samples in the discovery cohort. The red line represents the fitted linear regression, with the shaded area indicating the 95% confidence interval.

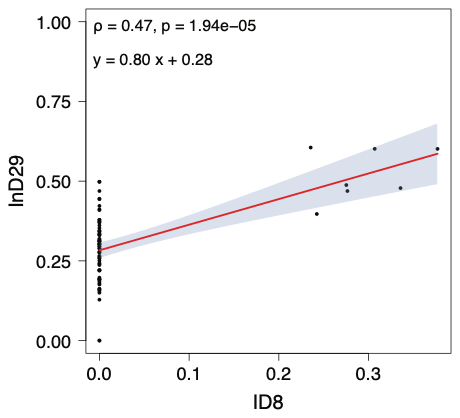

**Supplementary Figure 7. Distribution of InDel distances to SV breakpoints across SV classes.** Density plots showing the distribution of distances between somatic InDels and the nearest SV breakpoint within a 500 bp window, stratified by SV class: deletions (DEL), duplications (DUP), inversions (INV), and translocations (TRA). The dashed vertical line indicates the 100 bp threshold used to define proximal InDels.

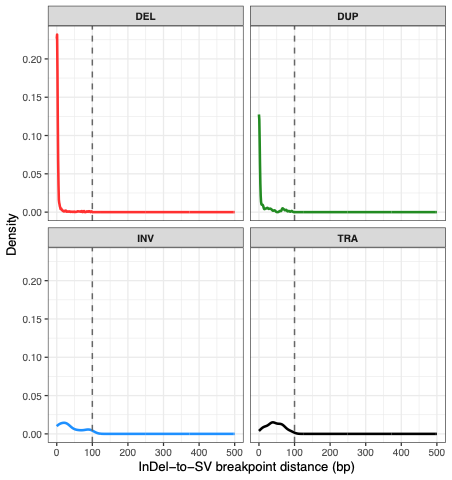

**Supplementary Figure 8. Expression of the 50 most highly expressed genes in MM.** Boxplots showing the distribution of gene expression levels across NDMM (blue) and RRMM (red) samples from the CoMMpass cohort. Genes were ranked by mean expression in NDMM and displayed in ascending order, with the same gene order maintained in RRMM. Expression levels are presented as log(TPM+1). Selected genes of biological relevance, including TXNDC5 (bold red), are annotated along the x-axis.

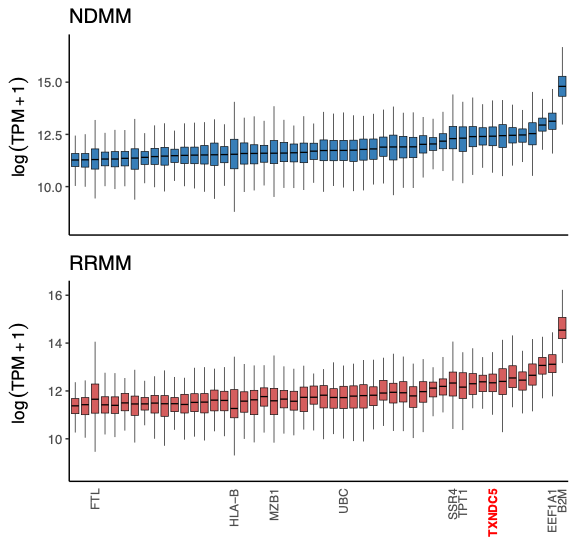

**Supplementary Figure 9. Evaluation of de novo InDel mutational signature extraction.** Reconstruction accuracy and stability metrics across solutions containing 1–5 signatures for the validation cohort (n = 421 samples). A five-signature solution was selected based on the balance between reconstruction accuracy and clustering stability.

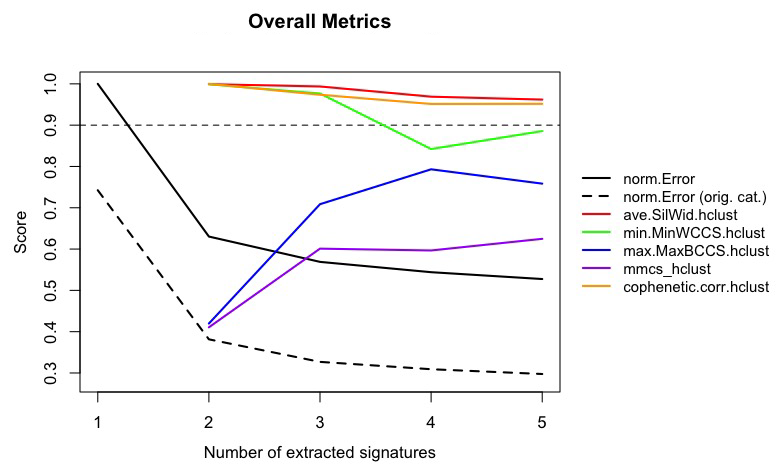

**Supplementary Figure 10. De novo extraction and deconvolution of InDel mutational signatures in the validation. cohort.** Mutational profiles of the five de novo InDel signatures (SIG1–SIG5) extracted from the validation cohort (n = 421 samples) using NMF and the 89-channel InDel classification framework. Bar heights represent the relative contribution of each mutational channel to the corresponding signature.

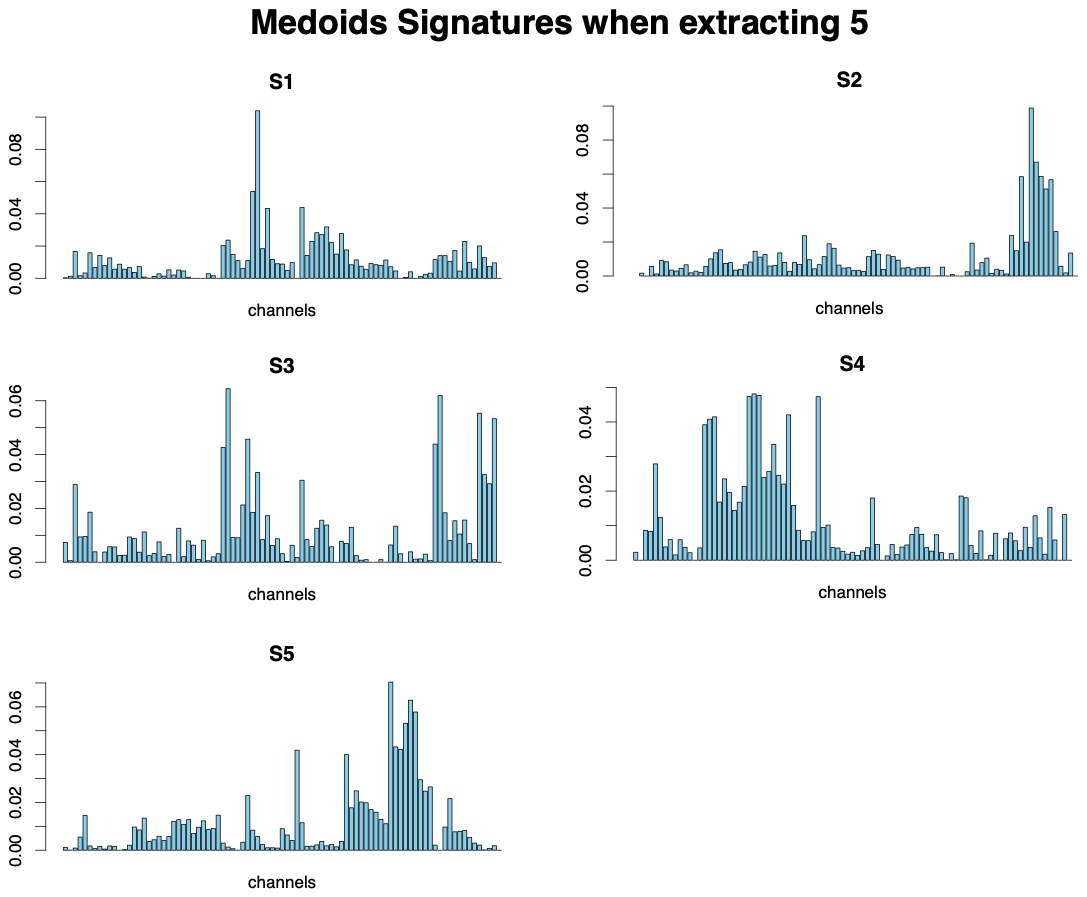

**Supplementary Figure 11.** Scatterplot illustrating the relationship between InD9a and SBS2/SBS13 mutational activity, in the NDMM samples part of the validation cohort (n = 421).

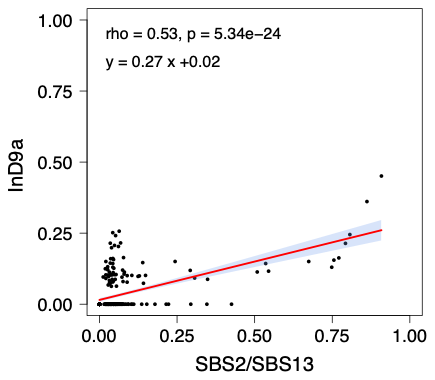

**Supplementary Figure 12.** 89-channel mutational profiles of 3,137 InDels located near SV breakpoints (±100 bp) in the validation cohort (n = 421).

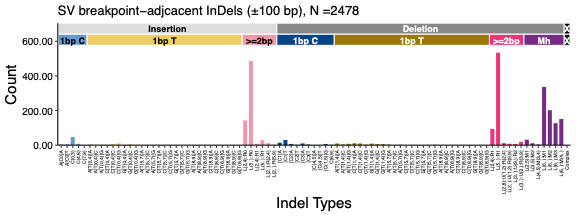

**Supplementary Figure 13.** SBS 96-mutational profile of MMRd+ case MM165_BL as well as its mutational signature contribution using *mmsig* fitting of SBS6 (MMRd).

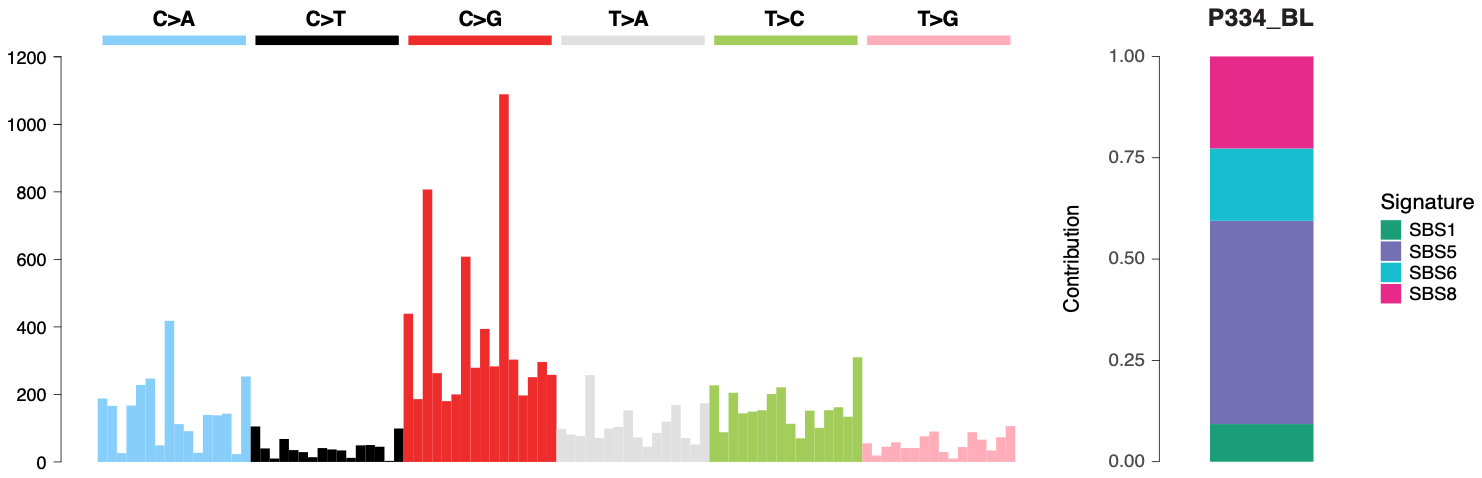

**Supplementary Figure 14. Epigenetic silencing of *MSH2* in 0661-1097_42188187_PDX_3-48_DNA_S1_001.** Representative genomic view of CpG methylation across the MSH2 promoter region in an MMRd+ PDX model. Individual sequencing reads are shown with methylated (red) and unmethylated (blue) CpG sites. CpG positions and corresponding methylation percentages are indicated below, alongside CpG sites previously investigated in neurofibromatosis type 1 (NF1) and extramammary Paget’s disease (1, 2).

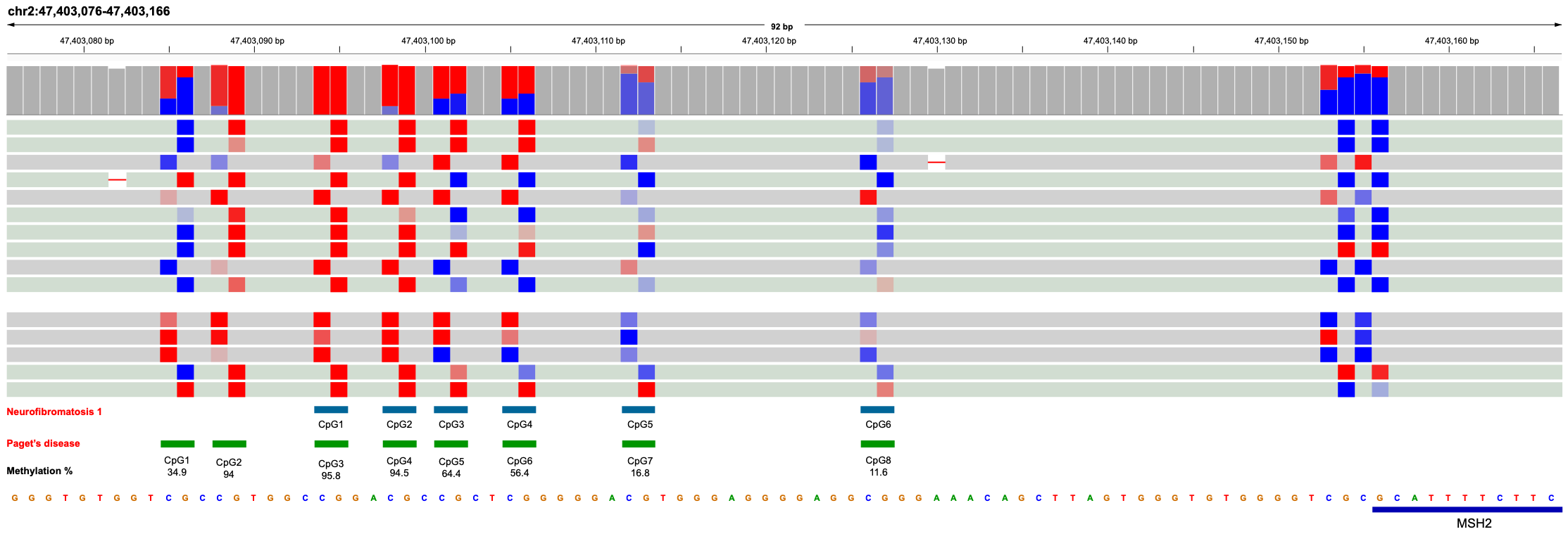
